# Height-induced postural threat selectively facilitates spinal reflex in the tibialis anterior muscle during quiet standing

**DOI:** 10.64898/2026.08.04.742625

**Authors:** Ryogo Takahashi, Naotsugu Kaneko, Keiichi Ishikawa, Kazuyuki Sato, Yume Mashiki, Kimitaka Nakazawa

## Abstract

Long-latency stretch reflex and corticospinal excitability in the tibialis anterior muscle (TA) are facilitated when balance is threatened, even without background TA activity, suggesting supraspinal modulation as preparatory tuning for ankle stabilization. However, it remains unclear whether such tuning is evident at the spinal level and specific to the TA among lower-limb muscles. We therefore examined the effects of height-induced postural threat on multi-segmental monosynaptic spinal reflexes (MMR) in lower-limb muscles during quiet standing. Seventeen healthy young males performed 90-s standing tasks under three postural threat conditions, created by combining real and virtual reality (VR) heights: (1) Low-threat (real ground & VR ground), (2) Medium-threat (real table & VR ground), and (3) High-threat (real table & VR bridge). During each condition, transcutaneous spinal cord stimulation (tSCS) was applied to the lumbar spine to elicit MMR in lower-limb muscles. Electromyograms (EMG) were recorded from six muscles of the right leg: vastus medialis (VM), biceps femoris (BF), TA, soleus (SOL), medial (MG), and lateral gastrocnemius (LG). MMR excitability was quantified as peak-to-peak EMG amplitude. Fear ratings and electrodermal activity were higher in High-threat than Low-threat (all *p* < 0.05), confirming successful threat induction. Peak-to-peak EMG amplitude in the TA was significantly higher in High-threat than Low-threat (17.1% increase, *p* = 0.0393), whereas background TA activity remained absent across conditions. These results indicate that TA has unique function to facilitate spinal excitability as a preparatory tuning for ankle stabilization.

**Key points:**

- Previous studies have shown the supraspinal facilitation of the tibialis anterior muscle without background muscle activation as a preparatory tuning for ankle stabilization.
- To test the hypothesis that such tuning is also evident at the spinal level and specific to the tibialis anterior muscle, this study examined whether height-induced postural threat modulates multi-segmental monosynaptic reflex excitability in lower-limb muscles using transcutaneous spinal cord stimulation.
- Electrodermal activity and fear ratings increased under height-induced postural threat, confirming the successful induction of postural threat.
- Under height-induced postural threat, the multi-segmental monosynaptic reflex was selectively facilitated in the tibialis anterior muscle, while its background activity remained absent.
- Our findings demonstrate selective facilitation of spinal excitability in the tibialis anterior muscle, which may serve as preparatory tuning for ankle stabilization under threat to balance.

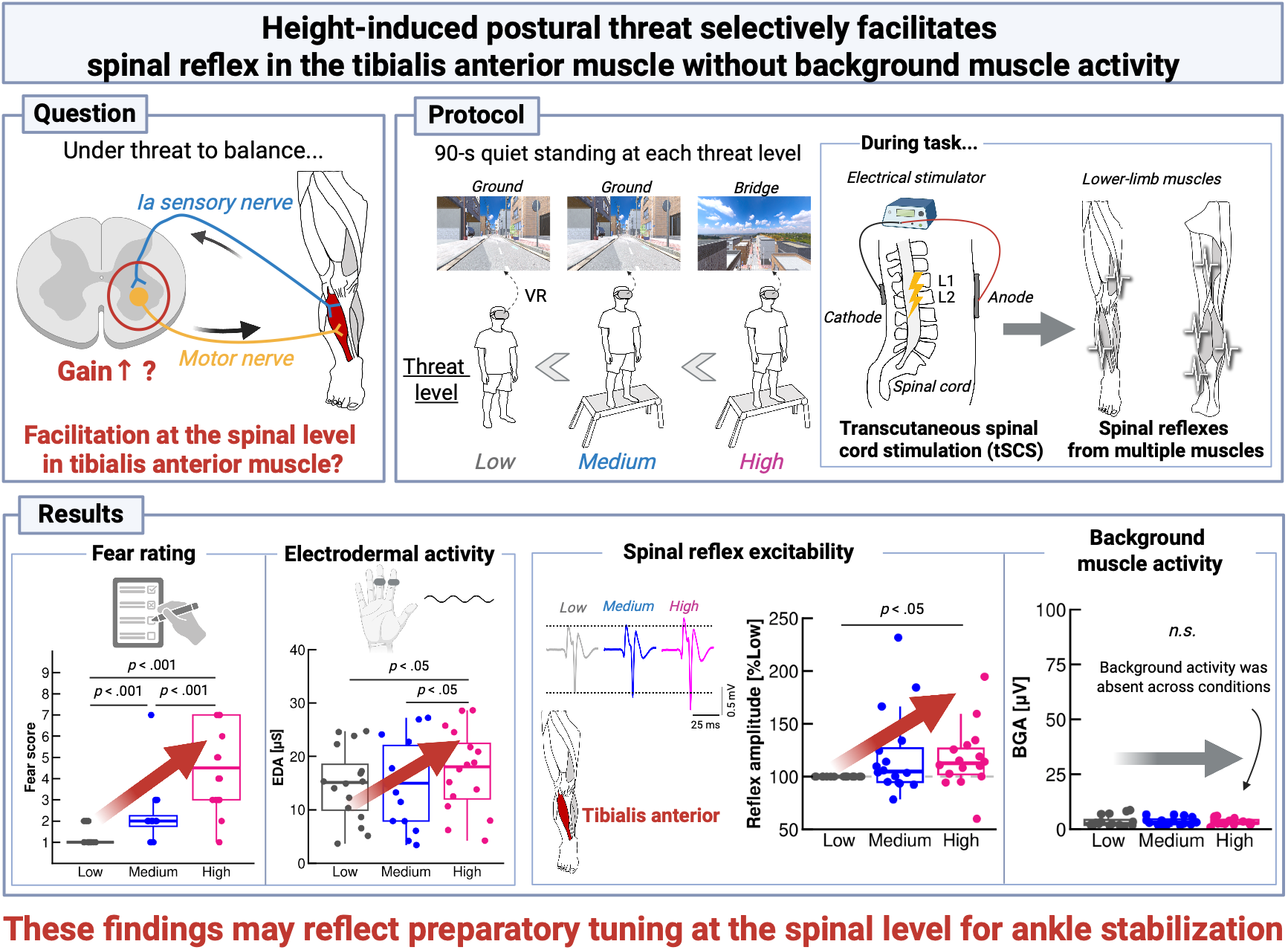

**Abstract figure legend:** When balance is threatened, corticospinal excitability and long-latency stretch reflex in the tibialis anterior muscle (TA) are facilitated even in the absence of background TA activity, suggesting supraspinal preparatory tuning for ankle stabilization. This study tested the hypothesis that such facilitation is also expressed at the spinal level and is specific to the TA. Participants completed 90-s quiet standing trials under three different height-induced postural threat conditions. During each trial, transcutaneous spinal cord stimulation was delivered over the lumbar spine to elicit multi-segmental monosynaptic reflexes (MMR) in multiple lower-limb muscles. High-threat condition increased fear ratings and electrodermal activity, indicating successful threat induction. Moreover, MMR excitability was selectively increased in the TA under High-threat condition despite the absence of background TA activity. These findings suggest that spinal facilitation is selectively expressed in the TA and may reflect preparatory tuning for ankle stabilization under threat to balance.

## Introduction

Bipedal upright standing and walking are mechanically unstable, characterized by a high centre of mass and small base of support (Winter, 1995). Such conditions can be interpreted as posing a potential threat to balance, as even a small perturbation may result in a loss of postural stability (Horak & Nashner, 1986). When balance is disturbed, the nervous system must therefore rapidly stabilize lower-limb joints to prevent a fall (Horak & Nashner, 1986). Ankle plays critical roles in stabilizing posture as it constitutes the primary mechanical interface between the body and the ground (Winter, 1995; Gatev *et al*., 1999).

Thus far, accumulating evidence indicates that motor pathway excitability related to tibialis anterior muscle (TA), ankle flexor, is enhanced under conditions in which balance is threatened (Christensen *et al*., 2001; Nakazawa *et al*., 2003, 2004, 2009; Fujio *et al*., 2016). For example, Christensen et al. showed the facilitation of long-latency stretch reflex and corticospinal excitability in the TA during the stance phase of gait, when ankle stabilization is particularly important (Christensen *et al*., 2001). Notably, these facilitations have been observed without background activity, suggesting that the long-latency stretch reflex and corticospinal excitability in the TA are modulated in preparation for ankle stabilization (Christensen *et al*., 2001). Consistent with this idea, facilitation of TA-related responses in the absence of background activity has been reported under several conditions in which balance is threatened. TA long-latency stretch reflex were facilitated during sudden surface drops while walking (Nakazawa *et al*., 2004) and when perturbations occurred without temporal predictability during standing (Fujio *et al*., 2016). Increased TA long-latency stretch reflex was also observed during sudden surface drops while standing (Nakazawa *et al*., 2009). Nakazawa et al. (2003) reported increases in both TA long-latency stretch reflex and corticospinal excitability during standing compared with the supine position, whereas comparable facilitation was not evident in ankle extensor, such as the soleus muscle (SOL). These findings suggest that the TA preparatory tuning for ankle stabilization under threat to balance is implemented through supraspinal mechanisms.

On the other hand, two questions remain unresolved regarding the physiological role of the TA under threat to balance: (1) whether preparatory tuning of the TA is also evident at the spinal level, and (2) whether this tuning is specific to the TA among lower-limb muscles. For the first question, the facilitation in TA long-latency stretch reflex and corticospinal excitability described above suggests the involvement of supraspinal modulation as a form of preparatory tuning for ankle stabilization under threat to balance. Recently, Unger et al. demonstrated that the TA H-reflex was facilitated during standing compared with the supine position when the TA was voluntarily activated (Unger *et al*., 2019). If a similar facilitation were observed during quiet standing, in which balance is threatened despite the absence of TA background activity, it would suggest that the TA prepares for a potential future loss of balance by subliminally increasing excitability at the spinal level. For the second question, threat to balance may enhance physiological arousal (Adkin & Carpenter, 2018, for a review). Such arousal-related changes, inducing noradrenergic and serotonergic pathways, could influence spinal excitability (Thorstensen *et al*., 2024). Therefore, it remains possible that threat-related changes in spinal excitability reflect generalized arousal-related modulation rather than TA-specific tuning. Therefore, simultaneous assessment of multiple lower-limb muscles is necessary to determine whether preparatory tuning under threat to balance is specific to the TA at the spinal level.

While H-reflex is well established method to evaluate excitability of spinal reflex pathway, it is difficult to elicit reliable H-reflex from the TA, particularly under resting states without muscle activation (Zehr, 2002; Unger *et al*., 2019). This poses a significant limitation as the TA is normally inactivated in young individuals during quiet standing (Masani *et al*., 2003), making it challenging to assess H-reflex excitability in the TA under quiet standing. In addition, H-reflex cannot be comprehensively elicited from multiple lower-limb muscles. Transcutaneous spinal cord stimulation (tSCS) is a non-invasive technique designed to overcome the technical limitations of the H-reflex (Minassian *et al*., 2007; Courtine *et al*., 2007). This method stimulates the dorsal roots of the spinal cord through the skin, enabling the elicitation of multi-segmental monosynaptic spinal reflexes (MMR) from multiple lower-limb muscles. MMR shares characteristics with the H-reflex (Courtine *et al*., 2007); specifically, it exhibits a short latency and is depressed by both Achilles tendon vibration and a 50 ms pre-stimulus.

The present study aims to investigate the effects of threat to balance on MMR excitability in lower-limb muscles using tSCS. We adopted the height-induced postural threat paradigm, in which participants stand quietly on platforms of different heights. This paradigm is a well-established method for manipulating the level of threat to balance by inducing threat-related emotional responses (e.g., unpleasant and aroused states) in participants (Adkin & Carpenter, 2018, for a review). We hypothesized that height-induced postural threat would increase TA MMR excitability without a concomitant increase in background muscle activity, suggesting modulation of spinal excitability as a preparatory tuning. By simultaneously recording MMR from multiple lower-limb muscles, we further examined whether this modulation reflected a generalized increase in spinal excitability across the lower limbs or muscle-dependent tuning specific to the TA. Clarifying this issue provides fundamental insight into the unique physiological role of the TA under threat to balance.

## Methods

### Participants

Prior to the experiment, we conducted a power analysis to determine the required sample size using G*Power (ver. 3.1). Specifically, the pilot experiment with 11 participants was conducted, and then the effect size of MMR amplitude of TA, as described in the Data analysis section, was calculated to estimate the required sample size (effect size (*η_p_*_2_): 0.183; *α* level: 0.05; power [1-*β* error probability]: 0.95). Since the data were originally non-parametric, we increased the estimated sample size by 15%, resulting in a required sample size of 16.1 participants (Ledolter & Kardon, 2020). Finally, seventeen healthy male volunteers with no history of mental or physical disorders were recruited for the study. The mean age, weight, and height of the participants with standard deviation (SD) were 25.2 ± 4.53 years, 66.1 ± 8.00 kg, and 174.1 ± 6.12 cm, respectively. Interoception, the process of monitoring internal bodily states (Craig, 2002), is associated with emotional response (Prentice *et al*., 2022) and posture (Dohata *et al*., 2025). Because interoception in females is influenced by the menstrual cycle (Prentice *et al*., 2022), we selectively recruited male participants to eliminate this potential confounding factor. All participants provided written informed consent to participate in the present study, and the experimental procedures were approved by the Institutional Review Board (IRB) of the University of Tokyo (Number: 792). This study was performed in accordance with the Declaration of Helsinki (1964).

### Data collection

Electrodermal activity (EDA) and Electromyogram (EMG) data were collected using two bipolar Ag/AgCl surface electrodes (Vitrode F-150S, Nihon Kohden, Tokyo, Japan) on the skin (Fig. 1A).

**Figure 1.**
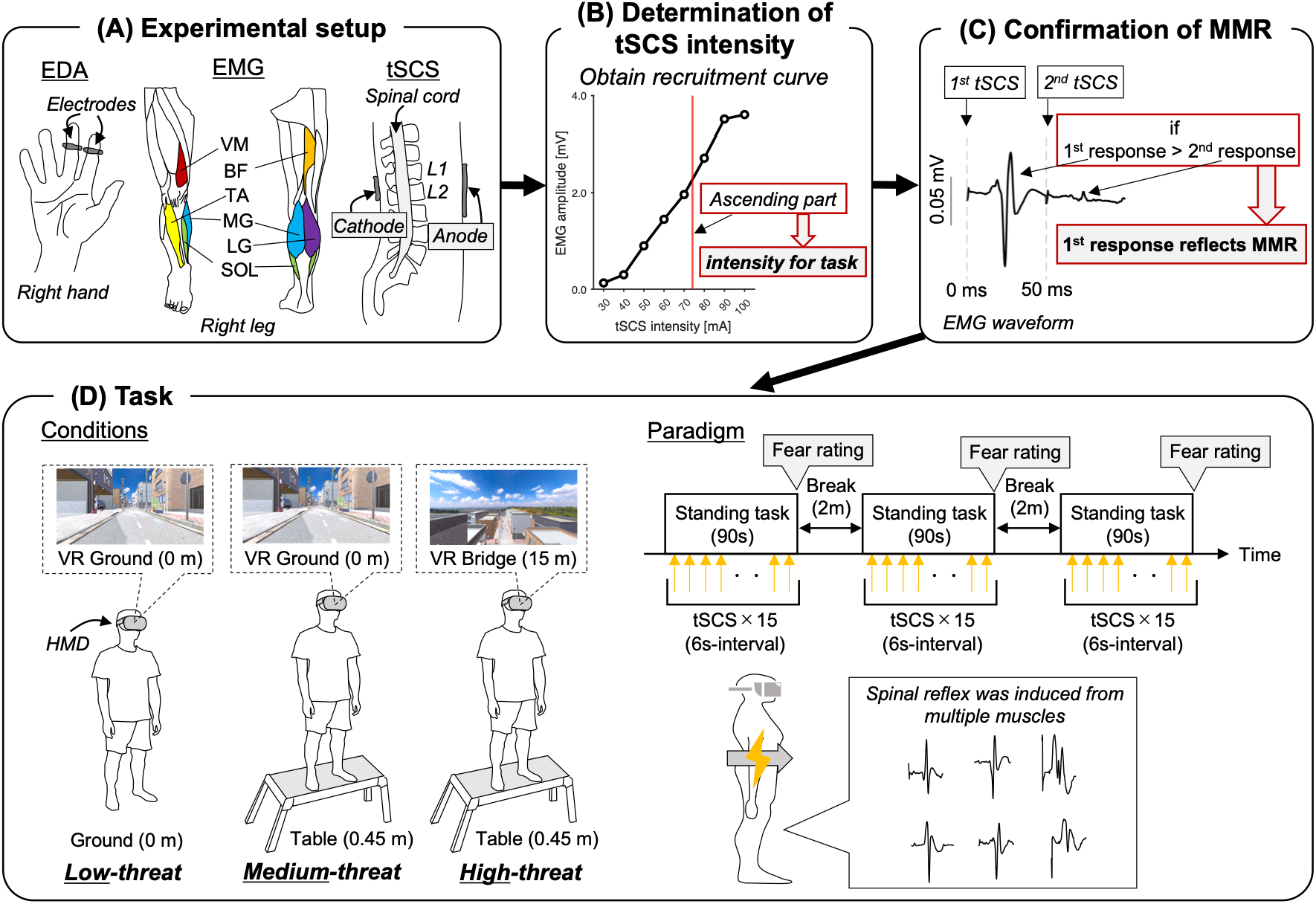
(A) Experimental setup for EDA, EMG, and tSCS. (B) Determination of tSCS intensity. Recruitment curve (input–output curve) was obtained from each muscle, and a tSCS intensity corresponding to the ascending part of the recruitment curves was used for the subsequent tasks. (C) Double-pulse tSCS was applied to determine whether tSCS evokes MMR via activation of sensory nerves rather than motor nerves. If the second EMG response was attenuated relative to the first, the first response was considered to represent the spinal reflex. (D) Experimental conditions and task paradigm. Three threat levels were established as experimental conditions by manipulating real and VR heights. In each condition, tSCS was applied 15 times during quiet standing. At the end of each task, participants reported their subjective fear level. A two-minute rest period was provided between conditions.

#### EDA recordings

EDA data, an index of sympathetic activity, were recorded from two electrodes placed on the middle and ring fingers of the non-dominant hand, and then amplified (×100) using a dedicated unit (AP-U030, Miyuki Giken, Tokyo, Japan). Finally, EDA data were recorded at a sampling rate of 1000 Hz using analog-to-digital (A/D) converters (MP208, Miyuki Giken, Tokyo, Japan).

#### EMG recordings

EMG data were recorded from the vastus medialis (VM), biceps femoris (BF), tibialis anterior (TA), soleus (SOL), medial head (MG), and lateral head of gastrocnemius (LG) muscles in right leg (Fig. 1B) in line with the SENIAM recommendation. Two electrodes were placed over the muscle belly with a 1.5 cm separation between the centre of electrodes. EMG signals were bandpass-filtered (5−1000 Hz) and amplified (×1000) using a multichannel amplifier (MEG-6108, Nihon Kohden, Tokyo, Japan). Subsequently, EMG data were recorded at a sampling rate of 4000 Hz using analog-to-digital (A/D) converters (Powerlab C, AD Instruments, Castle Hill, Australia).

### Transcutaneous spinal cord stimulation (tSCS)

The anode electrode (100 × 75 mm) was placed on the abdomen just above the navel, and the cathode electrode (50 × 50 mm) was placed on the middle of the first and second lumbar spine processes (L_1_-L_2_) (Fig. 1A). Next, we determined the optimal position of the cathode electrode. Specifically, participants were asked to stand quietly on a firm ground, and tSCS was delivered with the cathode electrode placed at T_12_–L_1_, L_1_–L_2_, or L_2_–L_3_ by using an electrical stimulator (DS7A, Digitimer Ltd. Hertford, United Kingdom). The position that elicited the greatest tSCS-induced EMG responses across all recorded muscles was selected as the cathode placement site (Th_12_–L_1_ for N = 1, L_1_–L_2_ for N = 16). To determine the stimulus intensity for the task, the recruitment curve was obtained by progressively increasing the stimulus intensity from 30 mA to 100 mA in 10 mA increments (Fig. 1B). To maximize the sensitivity of spinal reflex modulation to threat effects, the stimulus intensity where the recruitment curves of all muscles were on the ascending part was used for the task (Kaneko *et al*., 2021). As a result, the stimulus intensities were ranged between 50–90 mA (mean ± SD = 74.5 ± 12.1 mA) (Fig. 2). Next, we applied double-pulse stimulations (inter-stimulus interval: 50 ms) to confirm that tSCS-induced EMG responses are induced by sensory nerve activation but not motor nerve activation (Courtine *et al*., 2007) (Fig. 1C). When the second response is attenuated compared to the first response, the first response is originated by sensory nerve activation and considered as spinal reflex (Courtine *et al*., 2007). The double-pulse stimulations were repeated for 8 times, and then the first and second responses were averaged for each.

**Figure 2.**
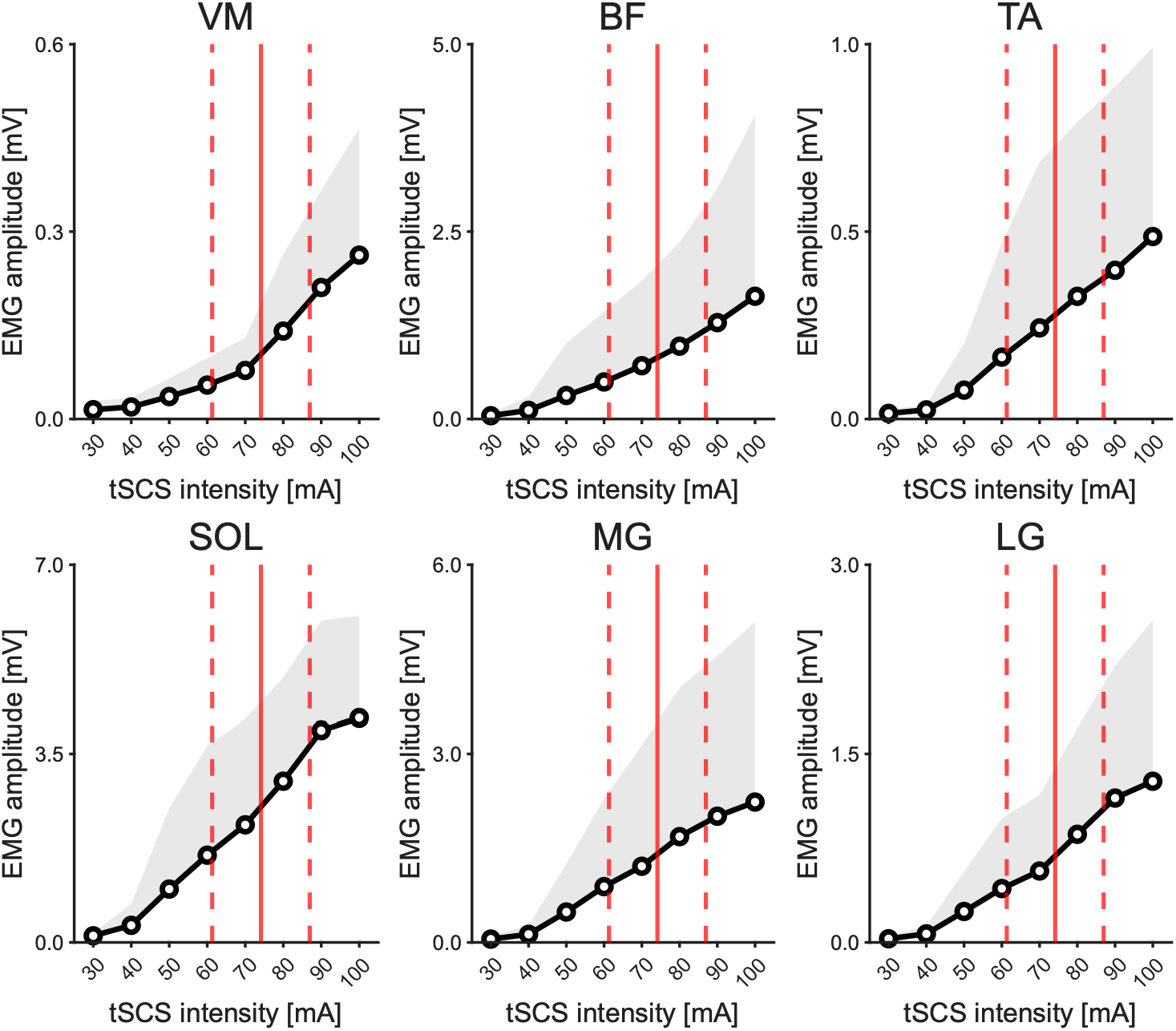
Recruitment curves of MMR. Black curves and shaded areas indicate averages and SDs of recruitment curves, respectively. Solid and dashed red lines shows averages and SDs of tSCS intensity used for the tasks, respectively.

### Task

In this study, a combined environment of reality and virtual reality (VR) was used to elicit an efficient threat for participants. A height environment induced by VR is well established tool to elicit autonomic, psychological, and postural changes similar to those induced by a real height environment (Cleworth *et al*., 2012). Accordingly, participants were asked to perform a 90-s quiet standing task with head-mounted display (Oculus 3, Meta, CA, USA) under different postural threat conditions: (1) Low-threat, (2) Medium-threat, and (3) High-threat conditions (Fig. 1D). Low-threat condition served as the baseline for calculating relative changes in physiological responses. Medium- and High-threat conditions were designed to examine whether different levels of threat influence physiological responses, consistent with a previous study demonstrating graded threat effects (Carpenter *et al*., 2001). In Low-threat condition, participants stood on the firm ground while experiencing VR in which they were standing on a street. In Medium-threat condition, participants stood on a table (height: 0.45 m) with their toes aligned with its edge while experiencing VR in which they were standing on a street. In High-threat condition, participants stood on a table (height: 0.45 m) with their toes aligned with its edge while experiencing VR in which they were standing on a bridge (height: 15 m). During each task, the single pulse tSCS was applied 15 times at 6 s intervals. After each task, participants rated how much fear they felt during the task using integers from 1 to 9, where 1 represented the lowest and 9 indicated the highest level of fear.

## Data analysis

All signals were analyzed using a custom-written script in MATLAB (2025b, MathWorks Inc., MA, USA). MMR amplitude (excitability) was defined by peak-to-peak EMG amplitude. Subsequently, the peak-to-peak EMG amplitude was normalized to the value obtained in the Low-threat condition. In addition, to assess background EMG activity (BGA), we calculated the mean amplitude of the rectified EMG signal within a 100 ms window preceding the tSCS. Trials in which the BGA exceeded + 2SD from the task mean were excluded from the analysis. The number of withdrawn trials was as follows: VM, 24/675 (15 trials × 3 conditions × 15 participants); BF, 25/720; TA, 30/720; SOL, 15/720; MG, 28/720; and LG, 31/720 (15 trials × 3 conditions × 16 participants each), demonstrating that fewer than 5% of trials were excluded for any muscle.

The raw EDA signals were low-pass filtered at 1 Hz using a fourth-order and zero-phase lag Butterworth filter. Subsequently, we calculated the mean EDA amplitude within a 100 ms window preceding the tSCS to quantify the sympathetic activity.

### Statistical analysis

All statistical comparisons were performed using the R software package (ver. 4.1.2). Shapiro-Wilk tests were conducted to confirm the normal distribution of the variables. Because certain variables showed non-normal distributions, we applied non-parametric statistical methods to all variables to maintain consistency in the statistical approach (Takahashi *et al*., 2025*b*). For the double pulse tSCS, we applied Wilcoxon singed-rank test to compare the first and second EMG responses. For task variables (BGA, MMR amplitude, mean EDA amplitude, and subjective fear rating), we conducted Friedman test to compare them across the three postural threat conditions. If Friedman test shows significant result, Wilcoxon signed-rank test was conducted as a post-hoc test. Additionally, to examine whether changes in MMR amplitude in response to postural threat were associated with subjective fear rating, mean EDA amplitude, and BGA, Kendall’s correlation coefficient (*τ*) was calculated between MMR amplitude and each of these variables. Prior to the correlation analysis, each variable was converted to a change ratio relative to the value in the Low-threat condition, calculated separately for the Medium- and High-threat conditions as follows:

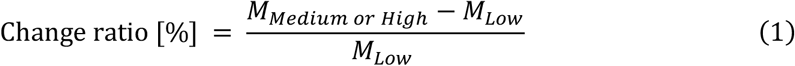

where *M_Medium or High_* is the value in the Medium- or High-threat condition, and *M*_*Low*_ is the value in the Low-threat condition. The change ratio for the fear rating was calculated only by subtracting the value in Medium- or High-threat condition from that in Low-threat condition (i.e., *M*_*Medium or High*_ – *M*_*Low*_). Correlation coefficients were then computed using the change ratios for each condition separately, as well as for the combined data from the Medium- and High-threat conditions. All *p*-values were corrected using the false discovery rate (FDR) correction, following the Benjamini-Hochberg procedure. The effect sizes for the Friedman test and Wilcoxon signed-rank test were calculated as Kendall’s W (*W*) standardized Z-statistic (*Z*), respectively. The significance level for all tests was set to *p* < 0.05.

## Results

### Double-pulse tSCS

Participants who did not exhibit even minimal attenuation of the second EMG response compared to the first were excluded from the EMG analysis. As a result, 15 participants were included for the VM, and 16 participants for the BF, TA, SOL, MG, and LG. Figure 3A indicates representative EMG waveforms of each muscle elicited by double-pulse tSCS. For all the muscles, the second response elicited by tSCS was significantly attenuated compared to the first response (Wilcoxon signed-rank test, all *p* < 0.001, Fig. 3B). The attenuation rate of each muscle was as follows (mean ± SD): VM, 53.13 ± 22.02%; BF, 78.02 ± 20.84%; TA, 81.24 ±13.88%; SOL, 92.53 ± 6.51%; MG, 84.23 ± 16.33%; LG, 91.70 ± 6.42%.

**Figure 3.**
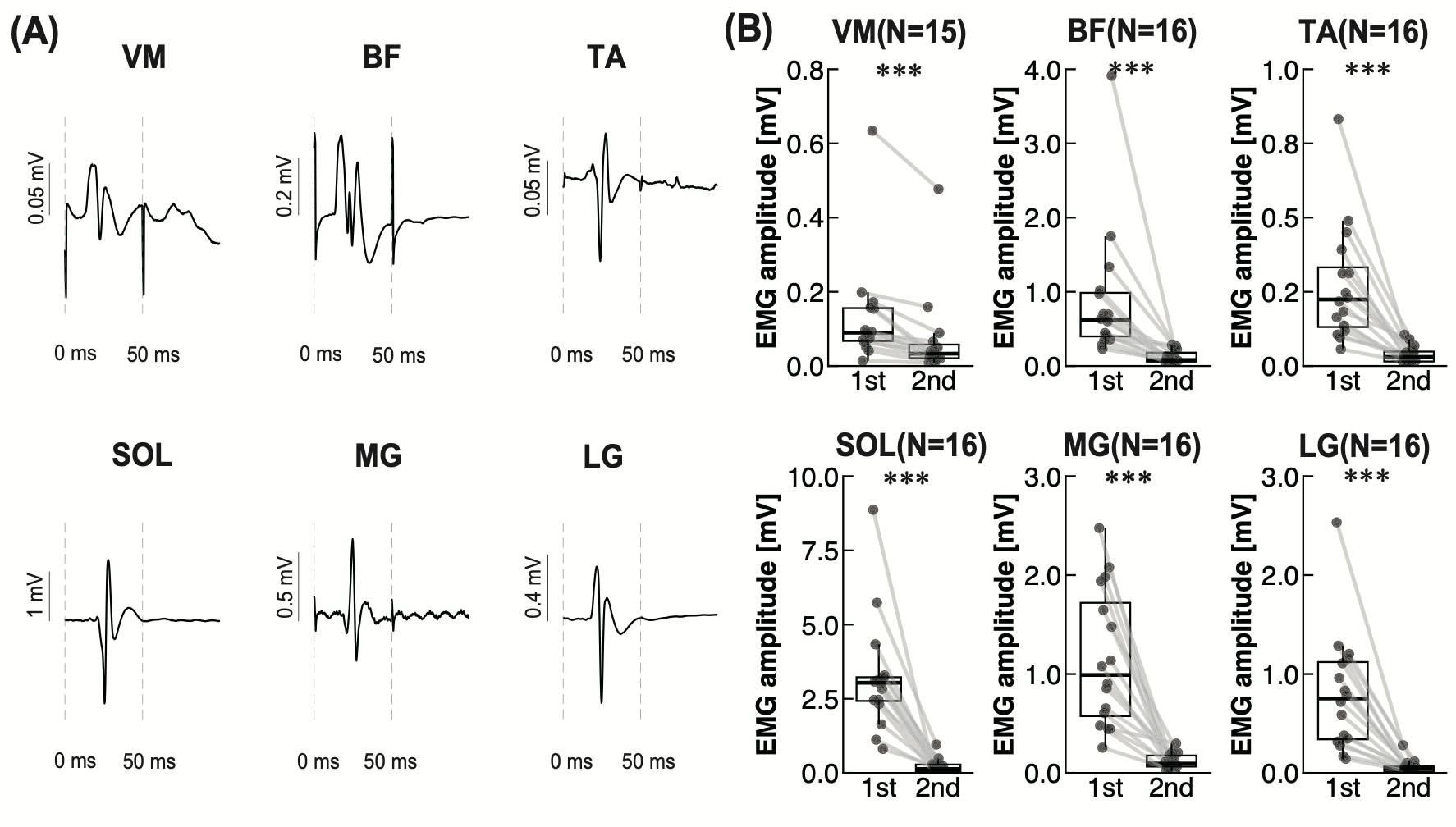
(A) Representative EMG waveforms induced by the double-pulse stimulation for one participant. They were computed by averaging EMG responses elicited by 8 trials. Vertical gray dot lines indicate each stimulus, first (0 ms) and second (50 ms) ones. (B) Group data for the amplitudes of first and second tSCS-evoked responses. The lines in the box plots indicate median values, and the ends of the boxes represent the 25th and 75th percentiles. The dots represent individual data points. \*\*\**p* < 0.001.

### Fear rating and sympathetic activity

One participant who showed no attenuation of second response induced by double-pulse tSCS in all the muscles was excluded from the analysis of fear rating and sympathetic activity; therefore, 16 participants were included in the analysis. Table 1 shows the mean values of all variables in each condition. Friedman test revealed a significant main effect of threat on fear rating (*p* < 0.001, Table 2). Post-hoc Wilcoxon signed-rank test showed that the value was significantly higher in High-threat condition than in Medium- and Low-threat conditions (Medium-threat: *p* < 0.001; Low-threat: *p* < 0.001, Table 3, Fig. 4A), and higher in Medium-threat conditions than in Low-threat condition (*p* < 0.001).

**Table 1.** Summary of the mean values (mean ± SD) of all variables in each condition.

|  | Conditions |  |  |
| --- | --- | --- | --- |
|  | Low-threat | Medium-threat | High-threat |
| *Emotional states and sympathetic activity* |  |  |  |
| Fear rating | 1.19±0.403 | 2.25±1.44 | 4.63±2.03 |
| EDA [μS] | 14.7±6.79 | 15.0±7.90 | 17.2±7.66 |
| *BGA* |  |  |  |
| VM [μV] | 3.00±3.65 | 2.88±1.39 | 3.30±1.76 |
| BF [μV] | 5.01±3.50 | 5.31±4.71 | 5.24±6.66 |
| TA [μV] | 3.53±2.59 | 3.50±1.83 | 3.29±1.49 |
| SOL [μV] | 25.5±12.8 | 26.8±12.4 | 25.6±10.3 |
| MG [μV] | 17.3±18.7 | 25.7±41.9 | 16.1±13.5 |
| LG [μV] | 6.29±5.06 | 6.34±4.18 | 7.03±5.82 |
| *MMR amplitude* |  |  |  |
| VM [mV] | 0.129±0.126 | 0.155±0.222 | 0.142±0.127 |
| BF [mV] | 1.15±1.16 | 1.35±1.65 | 1.18±1.49 |
| TA [mV] | 0.301±0.278 | 0.346±0.290 | 0.346±0.353 |
| SOL [mV] | 3.76±2.32 | 4.00±2.34 | 3.83±2.00 |
| MG [mV] | 1.33±0.759 | 1.54±0.742 | 1.40±0.706 |
| LG [mV] | 0.857±0.620 | 0.998±0.593 | 0.963±0.579 |

**Table 2.**
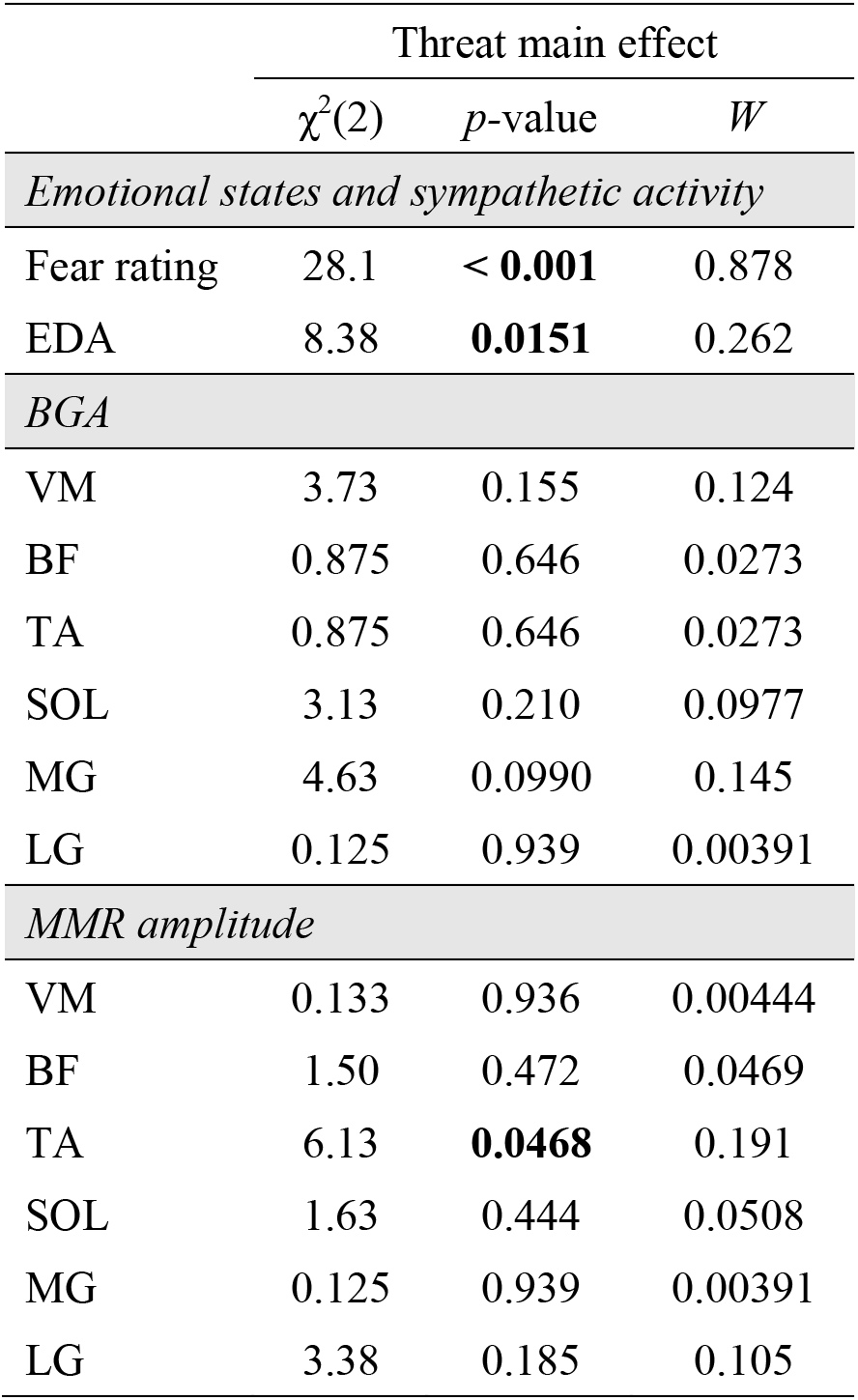
Summary of the Friedman test results for all variables. Significant *p*-values (< 0.05) are in boldface.

**Figure 4.**
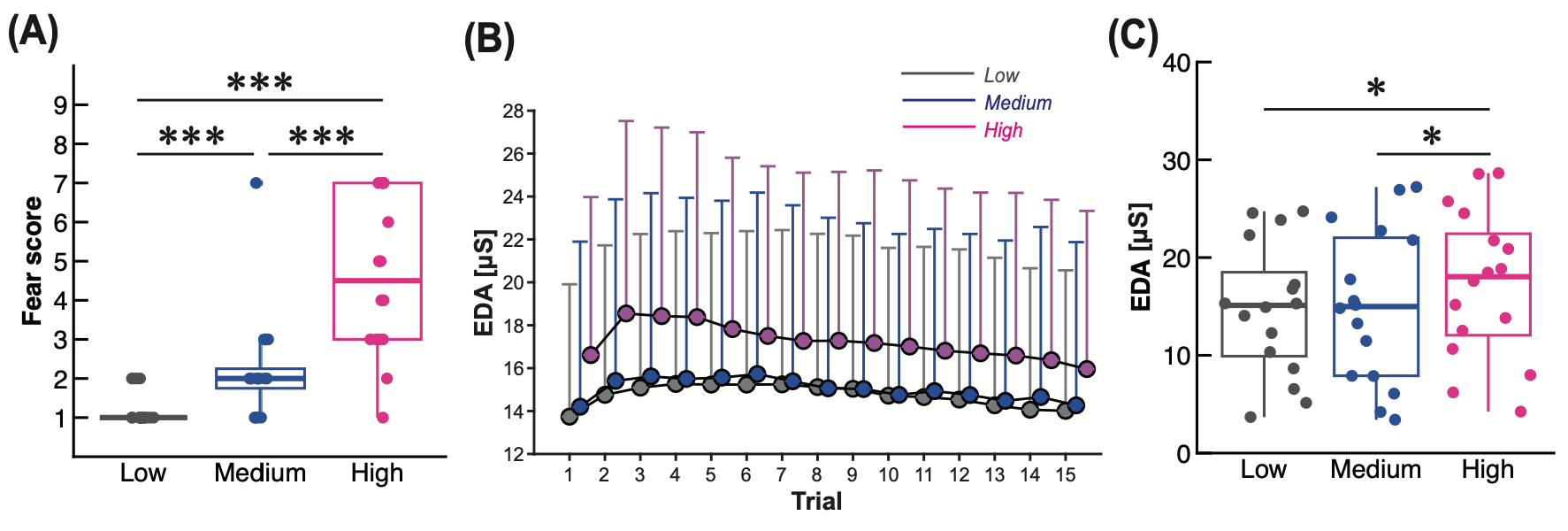
(A) Fear rating score. Low: Low-threat condition; Medium: Medium-threat condition; High: High-threat condition. The lines in the box plots indicate median values, and the ends of the boxes represent the 25^th^ and 75^th^ percentiles. The dots represent individual data points. \**p* < 0.05; \*\*\**p* < 0.001 (Wilcoxon signed-rank test). (B) Trial series of EDA amplitude. The circles indicate mean values, and the vertical lines represent SDs. (C) Mean EDA amplitude across trials.

**Table 3.**
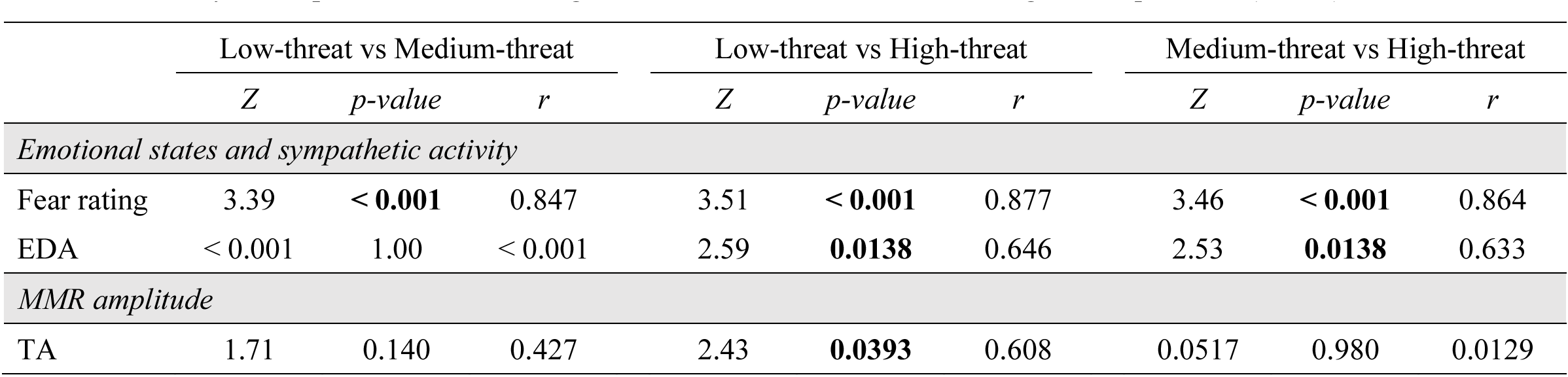
Summary of the post-hoc Wilcoxon signed-rank test results for variables. Significant *p*-values (< 0.05) are in boldface.

Figure 4B presents the trial series of EDA amplitude as sympathetic activity. Across all conditions, EDA amplitude reached its maximum during the first half of the trials and then gradually decreased thereafter. Regarding the averaged EDA amplitude across trials, a significant main effect of condition was observed (*p* = 0.0151, Table 2). High-threat condition showed significantly higher mean EDA amplitude than Medium- and Low-threat conditions (Medium-threat: *p* = 0.0138; Low-threat: *p* = 0.0138, Table 3, Fig. 4C).

### BGA

Friedman test revealed no significant main effect of threat for BGA across all muscles (all *p* > 0.05, Table 2, Fig. 5).

**Figure 5.**
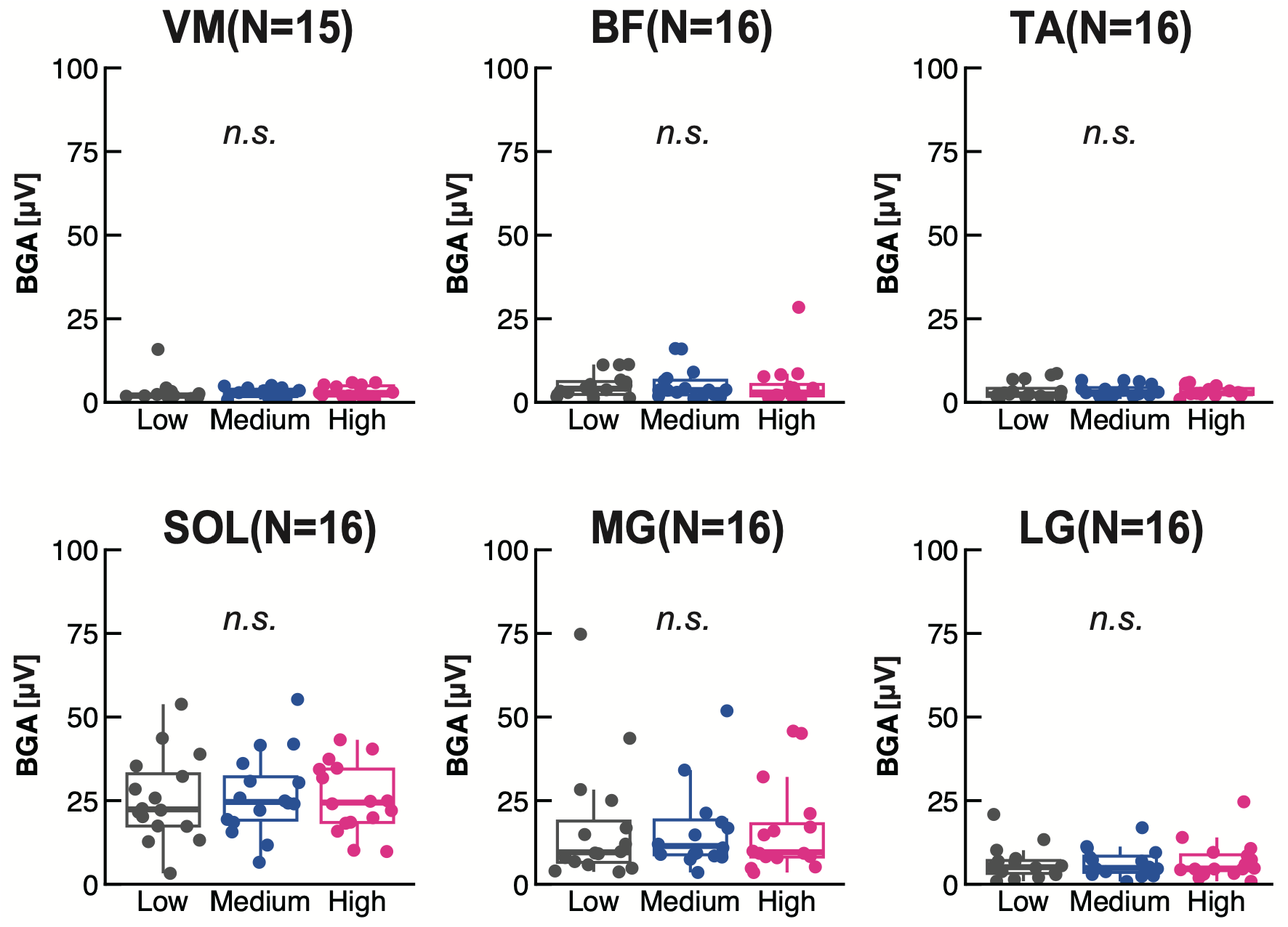
BGA of all muscles. Low: Low-threat condition; Medium: Medium-threat condition; High: High-threat condition. The lines in the box plots indicate median values, and the ends of the boxes represent the 25^th^ and 75^th^ percentiles. The dots represent individual data points. *n.s.* indicates no significant difference among conditions.

### MMR amplitude

Figure 6A shows representative waveforms of MMR from one participant. Friedman test revealed a significant main effect of threat on MMR amplitude in the TA (*p* = 0.0468, Table 2). Post-hoc Wilcoxon signed-rank test showed that the value was significantly higher in High-threat condition than in Low-threat condition (*p* = 0.0393, Table 3, Fig. 5B). On the other hand, no significant main effect of threat was observed for the other muscles (all *p* > 0.05, Table 2).

**Figure 6.**
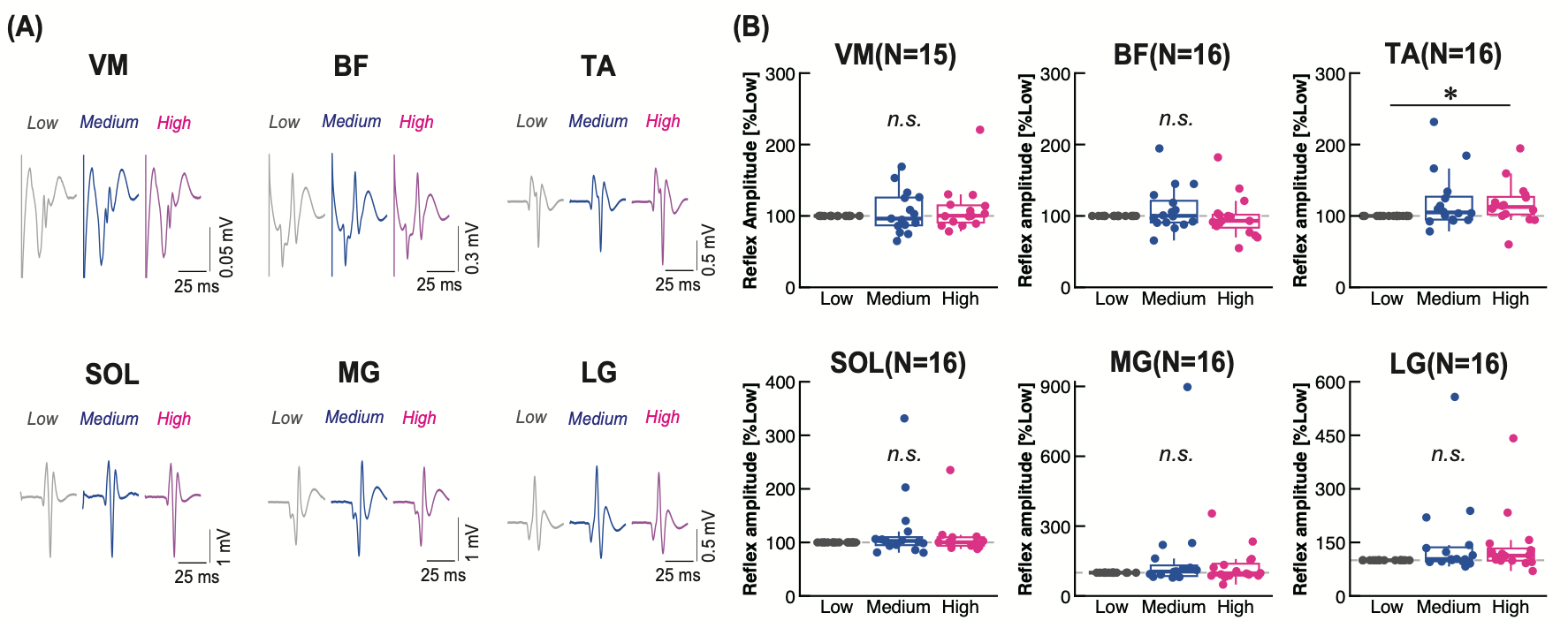
(A) Representative waveforms of MMR from one participant. Low: Low-threat condition; Medium: Medium-threat condition; High: High-threat condition. (B) MMR amplitude of all muscles. The values were normalized with Low-threat condition. The gray dashed line indicates 100%. The lines in the box plots indicate median values, and the ends of the boxes represent the 25^th^ and 75^th^ percentiles. The dots represent individual data points. \**p* < 0.05 (Wilcoxon signed-rank test). *n.s.* indicates no significant difference among conditions.

### Correlation analysis

Table 4 shows Kendall’s correlation coefficients between MMR amplitude and fear rating, EDA, and BGA. For all muscles, MMR amplitude was not significantly correlated with fear rating or EDA (all *p* > 0.05). In contrast, significant correlations were observed between MMR amplitude and BGA in the BF and MG (all *p* < 0.05), but not in the VM, TA, SOL, or LG (all *p* > 0.05).

**Table 4.**
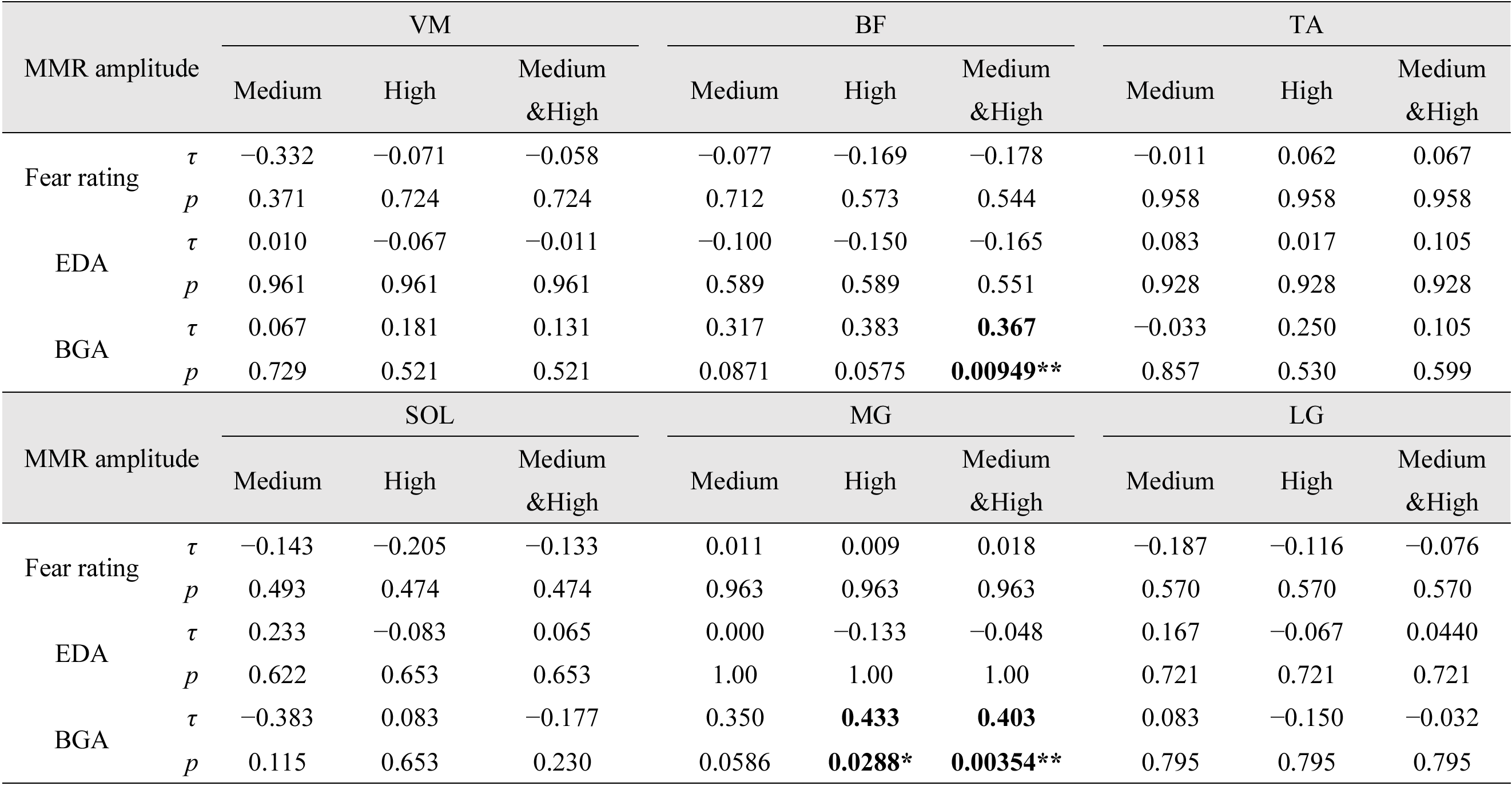
Kendall’s correlation coefficient (*τ*) between MMR amplitude and fear rating, EDA, and BGA for each muscle. Correlations were calculated using the change in each variable from the Low-threat condition, separately for the Medium- and High-threat conditions, and for their combined data. Correlation coefficients with significant *p*-values are in boldface. \**p* < 0.05; \*\**p* < 0.01.

## Discussion

This study investigated effects of height-induced postural threat on MMR excitability in lower-limb muscles during quiet standing. The double-pulse tSCS paradigm showed that the spinal reflex is successfully induced as the second responses were significantly attenuated compared with the first ones in the recorded muscles (Fig. 3B). Moreover, the subjective fear rating and EDA amplitude were significantly increased under height-induced postural threat, indicating that the threat to balance was successfully induced for the participants (Fig. 4). The main finding is that height-induced postural threat selectively increased MMR amplitude in the TA (Fig. 6) without the increase in BGA (Fig. 5), supporting our hypothesis. We discuss the functional meaning and neural mechanism of the series of results below.

### Functional meaning of facilitation in MMR excitability of TA

Previous studies have demonstrated that the long-latency stretch reflex and corticospinal excitability in the TA are facilitated under conditions in which balance is threatened, even in the absence of background TA activity (Christensen *et al*., 2001; Nakazawa *et al*., 2003, 2004, 2009; Fujio *et al*., 2016). These observations have been proposed to reflect preparatory tuning at the supraspinal level for ankle stabilization against potential balance loss (Christensen *et al*., 2001; Nakazawa *et al*., 2003, 2004, 2009; Fujio *et al*., 2016). The present study further revealed that height-induced postural threat facilitates MMR excitability in the TA without background activation, suggesting the existence of excitatory modulation at the spinal level. When standing balance is perturbed, afferent input from the muscle spindles elicits a monosynaptic stretch reflex via Ia fibers (i.e., short-latency stretch reflex) in the lower-limb muscles, enabling a rapid recovery from a potential fall (Nardone *et al*., 1990). It has been shown that greater excitability in the SOL H-reflex was associated with better recovery performance in response to a perturbation (Piirainen *et al*., 2013). Therefore, the enhancement of MMR excitability in the TA under height-induced postural threat may improve the early muscle response to perturbations. Given that the amplitude of the short-latency stretch reflex depends on the stretch velocity of the muscle spindles, increased MMR excitability in the TA under height-induced postural threat may particularly play role in the first balance recovery in response to a fast perturbation (Nakazawa *et al*., 2003; Horslen *et al*., 2018).

Although beyond the primary aim of this study, it is interesting to further discuss why the BGA in the TA did not significantly differ across conditions, given that some previous studies have reported TA activation under height-induced postural threat (Adkin & Carpenter, 2018, for a review). This discrepancy may be explained by the possibility that changes in the BGA in the TA are direction-dependent with respect to the perceived threat. When participants stand on a height platform with their toes along the edge of it, the body leans backward by accompanying increased TA and decreased ankle extensors (e.g., SOL and MG) activities (Carpenter *et al*., 2001). In contrast, when participants stand at the centre of an elevated platform, where the perceived threat is comparable in the anterior and posterior directions, ankle muscle activity shows minimal changes (Sibley *et al*., 2007). Similarly, in the present study, the High-threat condition required participants to stand on the narrow table and the virtual bridge with limited space in the anterior–posterior direction. In such a constrained environment, leaning the body forward or backward would not have been an appropriate strategy to prevent falling.

### Possible neural mechanism of facilitation of MMR excitability in TA

The observed facilitation of TA MMR excitability may be influenced by both afferent and descending pathways. Previous studies have shown that conditions in which balance is threatened enhance corticospinal excitability in the TA (Christensen *et al*., 2001; Nakazawa *et al*., 2003). In the context of height-induced postural threat, increased cortical involvement has also been demonstrated by studies using cortico-muscular and inter-muscular coherence (Zaback *et al*., 2022) as well as event-related cortical potentials (Zaback *et al*., 2023). Therefore, the cortical modulation may directly enhance intrinsic α-motoneuron (α-MN) excitability in the TA under height-induced postural threat, owing to their rich monosynaptic input from the motor cortex (Bawa *et al*., 2002; Eisner-Janowicz *et al*., 2023). Moreover, this cortical activation may lead to a reduction of presynaptic inhibition through the activation of interneurons that suppress presynaptic inhibitory mechanisms, thereby increasing Ia afferent input to α-MN (Meunier & Pierrot-Deseilligny, 1998). In addition to the cortical contribution, subcortical structures may also contribute to the MMR facilitation of the TA. It has been known that height-induced postural threat increases vestibulospinal and reticulospinal excitabilities (Horslen *et al*., 2014; Zaback *et al*., 2022, 2025). Regarding the afferent pathways, it is possible that increased sensitivity of muscle spindles enhances afferent input via Ia fibers, thereby increasing MMR excitability in the TA. Indeed, increased muscle spindle sensitivity has been well documented under height-induced postural threat (Horslen *et al*., 2013).

### Possible involvement of emotional systems

Height-induced postural threat is known to induce emotional changes associated with unpleasantness and enhanced physiological arousal (Adkin & Carpenter, 2018, for a review). The present study confirmed the lack of significant correlations between TA MMR excitability and fear ratings or EDA amplitude (Fig. 4). However, we cannot rule out the possibility that emotional factors influenced TA MMR excitability. This is because height-induced postural threat may also induce anxiety-related neural modulation, which may be partly dissociable from fear-related modulation (Pessoa, 2023). In the context of height-induced postural threat, vestibular-evoked short-latency EMG responses in the ankle extensors have been shown to be significantly associated with anxiety ratings, but not with fear ratings (Zaback *et al*., 2025). It is also possible that enhanced TA MMR excitability is related to arousal–valence dimensions, such as high arousal and unpleasant states, as well as to categorically defined emotions such as fear or anxiety. Although EDA amplitude reflects sympathetic activity as one component of physiological arousal, the absence of a significant correlation with EDA amplitude does not necessarily exclude the possibility that changes in arousal contributed to TA MMR excitability. Indeed, the effects of arousal and valence on postural control have been well documented in terms of ankle kinematics and kinetics (Horslen & Carpenter, 2011; Takahashi *et al*., 2024, 2025*a*) as well as neuromuscular activity in the ankle extensors (Takahashi *et al*., 2025*b*).

As possible emotion-related neural mechanisms, both descending and ascending circuits may be involved. It has been shown that recall of arousing episodic memories increases corticospinal excitability in the TA during sitting (Mashiki *et al*., 2025), suggesting emotion-dependent cortical input to spinal reflex circuit. Regarding subcortical regions, the vestibular nuclei and reticular formation have reciprocal neural connections with emotion-related regions, such as the amygdala (Staab *et al*., 2013). Moreover, high-arousal states may increase spinal excitability through serotonergic and noradrenergic projections (Thorstensen *et al*., 2024). In addition, unpleasant emotions increase muscle spindle sensitivity in the TA (Ackerley *et al*., 2017), possibly through γ-MN activation and enhanced muscle sympathetic outflow (Kamibayashi *et al*., 2009; Macefield & Knellwolf, 2018). Overall, the functional and anatomical interactions between motor and emotional systems may be associated with the facilitation of TA MMR excitability.

In addition, it would be interesting to investigate how height-induced postural threat influences MMR excitability in the lower-limb muscles of individuals with fear of falling. Comparing such responses with those observed in healthy individuals might help clarify the neurophysiological mechanisms underlying the association between fear of falling and increased fall risk (Maki *et al*., 1991).

### Limitations

We confirmed that the tSCS-induced EMG response is mediated by Ia afferent activation, as demonstrated by the double-pulse tSCS results (Fig. 3B). However, the second EMG response was not fully suppressed compared with the first response, suggesting that tSCS-induced EMG response may have contained direct motor nerve activation to some extent. Nevertheless, the ankle flexor and extensors, which were our primary focus, showed mean attenuation rates greater than 80%. In addition, because tSCS-induced EMG responses are expected to be influenced by spinal modulation, threat-dependent differences in these responses are likely to reflect modulation at the spinal level rather than changes in direct motor nerve activation alone.

### Conclusion

This study demonstrates that height-induced postural threat selectively facilitates TA MMR excitability without a concomitant increase in background EMG activity, whereas comparable modulation was not observed in other lower-limb muscles. These findings extend previous evidence of supraspinal facilitation of the TA as preparatory tuning for ankle stabilization by demonstrating that such tuning is selectively expressed in the TA at the spinal level under threat to balance.

## Funding

This work was supported by Japan Society for the Promotion of Science (JSPS) for Fellows Grant-in-Aid (KAKENHI) to R.T. [#24KJ0729; #26KJ0115], Descente and Ishimoto memorial foundation for the promotion of sports to R.T. [#2024], JSPS WAKATE to N.K. [#23K16745], JSPS Grant in Aid for Scientific Research (B) (General) to N.K. [#25K00127], Nakatani Foundation for Advancement of Measuring Technologies in Biomedical Engineering to N.K. [#2023], Tateishi Science Technology Foundation to N.K. [#2023A], Japan Science and Technology Agency (JST) PRESTO program to N.K. [#JPMJPR2415], and JST-MOONSHOT program to K.N. [#JPMJMS2012–2–2–2].

## Declaration of Competing Interest

The authors declare that the research was conducted in the absence of any commercial or financial relationships that could be construed as a potential conflict of interest.

## Author contributions

R.T. contributed to Conceptualization, Methodology, Data curation, Formal analysis, Investigation, Funding Acquisition, Visualization, and Writing - Original Draft. N.K. contributed to Supervision, Methodology, Investigation, Methodology, Funding Acquisition, and Writing - Review & Draft. K.I. contributed to Data curation and Writing - Review & Draft. K.S. contributed to Methodology and Writing - Review & Draft. Y.M. contributed to Data curation and Writing - Review & Draft. K.N. contributed to Conceptualization, Supervision, and Writing - Review & Draft. All authors contributed to the article and approved the submitted version.

## Data availability

The data presented in this manuscript were newly acquired for the present study. Because of data privacy concerns, they are not available to the community in open repositories. The datasets generated in this present study are available from the corresponding author upon reasonable request. In such cases, the reason for the data request and procedures for ensuring privacy will be reviewed and discussed.

## Notes

### Competing Interest Statement

The authors have declared no competing interest.

